# Machine Learning Models Identify Inhibitors of SARS-CoV-2

**DOI:** 10.1101/2020.06.16.154765

**Authors:** Victor O. Gawriljuk, Phyo Phyo Kyaw Zin, Daniel H. Foil, Jean Bernatchez, Sungjun Beck, Nathan Beutler, James Ricketts, Linlin Yang, Thomas Rogers, Ana C. Puhl, Kimberley M. Zorn, Thomas R. Lane, Andre S. Godoy, Glaucius Oliva, Jair L. Siqueira-Neto, Peter B. Madrid, Sean Ekins

## Abstract

With the ongoing SARS-CoV-2 pandemic there is an urgent need for the discovery of a treatment for the coronavirus disease (COVID-19). Drug repurposing is one of the most rapid strategies for addressing this need and numerous compounds have been selected for *in vitro* testing by several groups already. These have led to a growing database of molecules with *in vitro* activity against the virus. Machine learning models can assist drug discovery through prediction of the best compounds based on previously published data. Herein we have implemented several machine learning methods to develop predictive models from recent SARS-CoV-2 *in vitro* inhibition data and used them to prioritize additional FDA approved compounds for *in vitro* testing selected from our in-house compound library. From the compounds predicted with a Bayesian machine learning model, CPI1062 and CPI1155 showed antiviral activity in HeLa-ACE2 cell-based assays and represent potential repurposing opportunities for COVID-19. This approach can be greatly expanded to exhaustively virtually screen available molecules with predicted activity against this virus as well as a prioritization tool for SARS-CoV-2 antiviral drug discovery programs. The very latest model for SARS-CoV-2 is available at www.assaycentral.org.

## Introduction

In December 2019, several cases of pneumonia with unknown etiology started to arise in Wuhan, China. A new betacoronavirus was identified and named SARS-CoV-2 due to high similarity with previous SARS-CoV ^1,2^. This virus causes the disease which has been called COVID-19 ^3^.Since then, SARS-CoV-2 has rapidly spread worldwide prompting the World Health Organization to declare the outbreak a pandemic, with more than 1.5 million cases confirmed in less than 100 days.^4^ The high infection rate has also caused considerable stress on global healthcare systems leading to more than 400,000 deaths from COVID-19.

The SARS-CoV-2 pandemic started a worldwide effort to discover a treatment that could prevent further COVID-19 deaths and decrease the number and length of hospitalization^5^. Drug repurposing is one of the main strategies being used to accelerate this as most preclinical stages are removed and a promising drug could move directly into phase II clinical studies or beyond by using an approved, safe drug ^6,7^. So far, most SARS-CoV-2 inhibition studies rely on small to medium scale assays with high throughput screens (HTS) campaigns testing specific FDA-approved drugs and compounds that have previously shown inhibition against different betacoronaviruses or specific antiviral targets^8–16^.

Quantitative Structure Activity Relationship (QSAR) analyses from previous *in vitro* data has been widely used to assist drug discovery in both industry and academia^17^. In the past few years the rise of machine learning has also expanded to drug discovery, with different methods being implemented in a wide range of areas from predicting synthetic routes to biological activity^18,19^. Many examples show that prioritizing compounds from machine learning and QSAR models can increase the success rate and save resources^17^. Here we have implemented several machine learning methods to develop predictive models from recent SARS-CoV-2 *in vitro* inhibition data and used them to prioritize compounds for *in vitro* testing of different compound libraries. These efforts will add to the list of >200 drugs and vaccines under assessment elsewhere and which is continually growing ^20^.

## Materials and Methods

### Data Curation

Data from recent drug repurposing campaigns for SARS-CoV-2 were used to build a dataset from whole cell inhibition assays ^8,9,12,14,15^. In assays with several Multiplicity of Infection (MOI) the one closer to the whole dataset was chosen. In machine learning model generation, duplicate compounds with finite activities are averaged into a single entry. Due to the potential for diminished activity, when duplicate compounds were present, only the most active one was retained in the dataset. Additionally, compounds with ambiguous dose-response curves were discarded. Datasets were built with Molecular Notebook (Molecular Materials Informatics, Inc). In order to evaluate the model performance on an external testing set, a total of 30 molecules was collated from different studies^11,21–25^.

### Assay Central™

The Assay Central™ software (AC) has been previously described^19,26–34^. AC employs a series of rules for the detection of problem data for automated structure standardization to generate high-quality data sets and Bayesian machine learning models capable of predicting potential bioactivity for proposed compounds. AC was used to prepare and merge data sets, as well as generate Bayesian models using the ECFP6 descriptor and five-fold cross validation. During model generation, training compounds are standardized (i.e. salts were removed, corresponding acids neutralized), and thresholds for binary activity classification are applied to optimize internal five-fold cross validation metrics. For predictions, AC workflows assign a probability score and applicability score to prospective compounds according to a user-specified model, with prediction scores greater than 0.5 considered active.

### Additional Machine Learning Methods

Additional Machine learning algorithms including Bernoulli Naïve Bayes (bnb), AdaBoost Decision trees (ada), Random Forest (rf), support vector classification (svc), k-Nearest Neighbors (knn) and Deep Learning (DL) were also implemented with ECFP6 fingerprints and five-fold cross validation. Details for the development of these models was previously described in detail in our earlier articles ^28,32,35^. Bayesian models were also generated with Discovery Studio (Biovia, San Diego CA) using ECFP6 descriptors where the top and bottom scoring fingerprints were selected for qualitative comparison.

### Model Performance

Machine learning model performance was evaluated with different metrics: accuracy, recall, precision, specificity, F1-score, area under receiver operating characteristic curve, Cohen’s kappa, and the Matthews correlation coefficient. The statistics were calculated for both training data with five-fold cross validation, to evaluate training performance, as well as in external testing set, to evaluate model performance in predicting data outside the training set.

### Principal Component Analysis

Principal Component Analysis (PCA) was computed for both the SARS-CoV-2 data set as well as SARS-CoV-2 with different compound libraries to assess its chemical space. The scikit-learn^36^ (0.22.2) PCA algorithm was used to reduce feature dimensionality to three using different molecular descriptors (MW, MolLogP, NR, NArR, NRB, HBA, HBD) and also with EFCP6 fingerprints. Molecular descriptors and fingerprints were generated from the cheminformatics library RDkit (2020.03.1).

### Applicability and Reliability Domain Assessment

In order to check if it is valid to apply the model for compounds being predicted and how reliable the predictions are, an applicability and reliability domain assessment was performed. First, the compound applicability within the model is assessed comparing its similarity with the model’s data using both molecular and fingerprint descriptors. If the molecule satisfies both criteria it is considered within the applicability domain and goes to the reliability domain assessment.

The first criterion for the applicability assessment is determined based on whether it fits within the range of the key molecular descriptors of the training set (MW, MolLogP, NRB, TPSA, HBA, HBD). If at least four properties lie within the maximum and minimum values of the model’s data, the molecule is considered similar and goes to the next criterion. The second criterion relies on structural fragment-based similarity measured with Tanimoto coefficient using MACCS fingerprints. The similarity of the MACCS fingerprints for the query compound and all training data is computed using the Tanimoto score. Only 5% of the training set compounds that are most similar to the query compound is used for evaluation (i.e. if the training set has 100 molecules only 5 molecules with more similarity to the query compound are used for the next evaluation). If the Tanimoto score exceeds 0.5 against the 5% of the training set compounds, the model is considered to have enough structural fragments overlap with the query compound and thus the compound goes onto the reliability assessment.

The reliability domain assessment implements k-means clustering methods based on ECFC6 fingerprints to classify the predictions from very high to low reliability. The reliability class depends on four criteria: distance from the major central point of the training data, distance from the closest cluster, closest cluster density and closest cluster distance within the chemical space. Each criterion has different weights and scores, with the second and third having higher priority. If the compound scores 1 in each criterion it is classified as very highly reliable, if that is not the case only the two higher priority criteria are considered for the next classes. The compound is classified as highly reliable if scores a total of 2, moderately reliable if it scores between -1 and 2 or low reliability if it scores less than or equal to -1 in the two higher priority criteria. The scores for each criterion as well as its definition are extensively described in the Supplemental Methods.

### *In vitro* testing

Compounds were tested in a 10-point serial dilution experiment to determine the 50% inhibitory concentration (IC_50_) and 50% cytotoxicity concentration (CC_50_). 1,000 HeLa-ACE2 cells/well were added into 384-well plates with compounds in a volume of 25 nl. The final concentrations of compound ranged from 78nM-40µM. 4 h post seeding 500 pfu SARS-CoV-2 (Washington strain USA-WA1/2020), BEI Resources NR-52281 were added to each well at a MOI = 0.5. Twenty-four hours post infection cells were fixed with 4% formaldehyde solution. The cells were then treated with a Primary ab: human polyclonal plasma (COVID-19 patient); Secondary ab: goat anti-human IgG coupled with HRP. Images were acquired with ImageXpress MicroXL (bright field); Custom Module developed in MetaXpress was used for automated count of total cells and infected cells. Antiviral activity was assessed based on the infection ratio (number of infected cells/total number of cells) in comparison with the average infection ration of the untreated controls.

## Results

### Data Curation

*In vitro* SARS-CoV-2 data was initially collated from five drug repurposing studies leading to a data set of 63 molecules with mean activity of 15.94 ± 22.45 μM ^8,9,12,14,15^. The external testing set collated from different studies has 30 molecules and a mean activity of 34 ± 42 µM ^11,21–25^. Most assays were performed with different Vero cell lines, inhibition was measured with viral RNA quantification, cytopathogenic effects or immunofluorescence methods with MOI and incubation time varying from 0.01-0.05 and 24-72 hrs respectively (Figure S1). The threshold set for activity classification by the Bayesian model generated with AC was 6.65 µM, with a final ratio of 52% actives in the training set and 37% in the external test set. The molecules in both training and test set are available in the supplemental data.

### Machine Learning Models

Machine learning models were developed with AC as well as several other methods available to us. This five-fold cross validation comparison shows the different prediction statistics for all machine learning algorithms implemented with the training data only (Table 1). AC outperformed all of them at the threshold of 6.65 μM with Rf coming the closest. These models were chosen for further external testing predictions.

**Table 1.** Five-fold cross validation statistics for all SARS-CoV-2 machine learning models implemented using ECFP6 fingerprints.

|  | ACC | AUC | CK | MCC | Pr | Recall | Sp | F1 |
| --- | --- | --- | --- | --- | --- | --- | --- | --- |
| **AC** | 0.81 | 0.78 | 0.62 | 0.64 | 0.78 | 0.88 | 0.73 | 0.83 |
| **rf** | 0.75 | 0.74 | 0.49 | 0.5 | 0.73 | 0.82 | 0.67 | 0.77 |
| **knn** | 0.71 | 0.71 | 0.43 | 0.42 | 0.71 | 0.76 | 0.67 | 0.74 |
| **svc** | 0.7 | 0.69 | 0.39 | 0.4 | 0.68 | 0.79 | 0.6 | 0.73 |
| **bnb** | 0.68 | 0.68 | 0.36 | 0.36 | 0.7 | 0.7 | 0.67 | 0.7 |
| **ada** | 0.64 | 0.63 | 0.27 | 0.26 | 0.65 | 0.67 | 0.6 | 0.66 |
| **DL** | 0.65 | 0.65 | 0.3 | 0.3 | 0.66 | 0.67 | 0.63 | 0.66 |

ACC: Accuracy, AUC: Area under curve, CK: Cohen's Kappa, MCC: Matthews correlation coefficient, Pr: Precision, Sp: Specificity, F1: F1 Score. bnb: Bernoulli Naïve Bayes, ada: AdaBoost Decision trees, rf: Random Forest, svc: support vector classification, knn: k-Nearest Neighbors and DL: Deep Learning (DL)

### External Validation

The performance of the machine learning models on the external testing data is shown in Table 2. The external validation was used to measure model performance in data from different studies outside the training set. svc and knn had slightly better statistics compared to all other models, with the best balance between recall and specificity.

**Table 2.** Prediction statistics with the external data for all SARS-CoV-2 machine learning models implemented

|  | ACC | AUC | CK | MCC | Pr | Recall | Sp | F1 |
| --- | --- | --- | --- | --- | --- | --- | --- | --- |
| **AC** | 0.62 | 0.58 | 0.17 | 0.17 | 0.50 | 0.40 | 0.76 | 0.44 |
| **rf** | 0.63 | 0.57 | 0.10 | 0.11 | 0.42 | 0.30 | 0.80 | 0.35 |
| **knn** | 0.67 | 0.6 | 0.21 | 0.21 | 0.50 | 0.40 | 0.80 | 0.44 |
| **svc** | 0.70 | 0.57 | 0.34 | 0.34 | 0.54 | 0.60 | 0.75 | 0.57 |
| **bnb** | 0.50 | 0.49 | -0.09 | -0.09 | 0.27 | 0.30 | 0.60 | 0.28 |
| **ada** | 0.53 | 0.49 | 0.00 | 0.00 | 0.33 | 0.40 | 0.60 | 0.36 |
| **DL** | 0.63 | 0.56 | 0.15 | 0.15 | 0.44 | 0.40 | 0.75 | 0.42 |

### Chemical Space

The PCA of the model training set alone shows that the SARS-CoV-2 chemical space is well distributed with active and inactive molecules well mixed when analyzed using either molecular and fingerprint descriptors. When compared with Prestwick Chemical Library (PwCL), a library of predominantly FDA approved drugs, the SARS-CoV-2 data lie within a big cluster with molecular descriptors and is more widely distributed when using the fingerprint descriptors.

### Applicability and Reliability Domain Assessment of External Test Set

The applicability and reliability domain assessment of the external test set was determined for each molecule as described in the methods to see how the test set compares with the training data. Molecules in the applicability domain are considered suitable for the model predictions due to similarity based on structural and molecular properties with the training data, whereas the reliability value is a measurement of how reliable the predictions are and uses different clustering metrics to determine its value.

From 30 molecules in the external test set, 22 were within the training data applicability domain and had their reliability value calculated. Most molecules that fell within the applicability domain had high or very high reliability values, with only 36% showing moderate reliability, so, most molecules obey the similarity criteria and are not far away from dense clusters. In comparison, with the Assay Central applicability score, which accounts only for structural similarity of the query compound with the training data, only 10 molecules were considered within the domain with a higher reliability, suggesting it is likely more conservative. Indeed, with the external test and training set PCA we can see that most molecules superimpose with few of them distant from each other (Figure S1). Therefore, similarity together with clustering methods are more suitable for applicability and reliability assessment compared with only structural similarity, as seen by the PCA.

### Prospective Prediction

A selection of FDA approved drugs available to us in our relatively small in-house compound collection of hundreds of molecules was scored with the AC Bayesian model. A selection of some of the best scoring molecules (Table 3) was used to identify and prioritize compounds for *in vitro* testing. AC Applicability score is the similarity of the compound with the training data, compounds are ranked by reliability which may provide some degree of confidence in these predictions.

**Table 3.** Prospective prediction compounds predicted and prioritized for testing.

| Name | Prediction Score | AC Applicability Score | Reliability |
| --- | --- | --- | --- |
| CPI1062 | 0.67 | 0.5 | High |
| CPI1066 | 0.62 | 0.38 | High |
| CPI1004 | 0.62 | 0.39 | High |
| CPI1012 | 0.70 | 0.70 | Moderate |
| CPI1155 | 0.70 | 0.40 | Moderate |
| CPI1175 | 0.65 | 0.41 | Moderate |
| CPI1153 | 0.7 | 0.7 | Low |

### *In vitro* Inhibition Assays of Predicted Compounds

Antiviral activity testing in the HeLa-ACE2 cells demonstrated that CPI1155 and CPI1062 have antiviral activity with IC_50_ values of 8.4µM and 540nM (Figure 2), respectively. The cell viability of these compounds was also tested, with both CC_50_ higher than 40 µM. Other compounds did not inhibit viral replication in HeLa cells or had appreciable cytotoxicity.

## Discussion

One of the challenges for addressing novel viral outbreaks is selection of drugs to test. Testing capacity, even for *in vitro* antiviral activities is likely to be low at the onset of an outbreak, making compound selection even more critical in this situation. In the case of SARS-CoV-2, the initial focus was on molecules that had previously shown activity against SARS or MERS ^37,38^. The training set for the current model is therefore not a random sampling of drug property space. When compared with the PwCL, a library of mostly FDA approved drugs, all molecules superimpose in the property space highlighting the model suitability for drug repurposing. Even with a relatively small training dataset the machine learning models evaluated have shown acceptable five-fold cross validation statistics, with almost all metrics greater than random and ROC >0.75 for AC (Table 1). When compared with different machine learning methods AC outperforms all of them in the SARS-CoV-2 training set, but this may be due to the threshold for all models being set as optimal for AC. However, choosing different values could imbalance the training set and remove important compounds from the active group.

More important than a good performance in the training set is the performance on external data, since most prospective predictions will occur for molecules outside training data. For external validation all models had intermediate performance, with ROC of 0.6. Taking into account the small number of molecules and that some test set molecules lie outside the applicability domain, the performance is acceptable. Different from the training set performance, svc had the highest overall score, predicting 60% of the active molecules despite its modest statistics in five-fold cross validation. The good performance of svc in predicting biological activity is in accordance with several studies that show good performance in different datasets ^28,32,35,39^. Therefore, the models described here are suitable for initial prospective predictions.

The applicability and reliability assessment shows that 73% of the test set molecules lie within the model applicability domain with high to moderate reliability, so poor performance in external validation occurs because there isn’t a clear boundary in the model’s feature space that can correctly classify external data. Increasing the number of molecules might include new features in both actives and inactive molecules which can increase model performance in both training and external data.

The training and test set described herein can be merged to increase data set size and applicability domain. The AC model with merged training and test data has slightly worse statistics (ACC: 0.76, AUC:0.79, CK: 0.53, MCC: 0.75, Pr: 0.76, Recall: 0.76, Sp: 0.77, F1: 0.76), but a higher applicability domain. The PCA confirms this wide chemical property space (Figure S1), the PCA of this updated model is much more balanced and broader than the previous one (Figure S2) versus Figure 1B. Without some form of external validation, we cannot assess how predictions of compounds outside the applicability domain perform, as model statistics were comparable it is expected that compounds outside this would obviously have unreliable predictions, however this may be offset by a higher domain which can increase reliability of some compounds.

**Figure 1.**
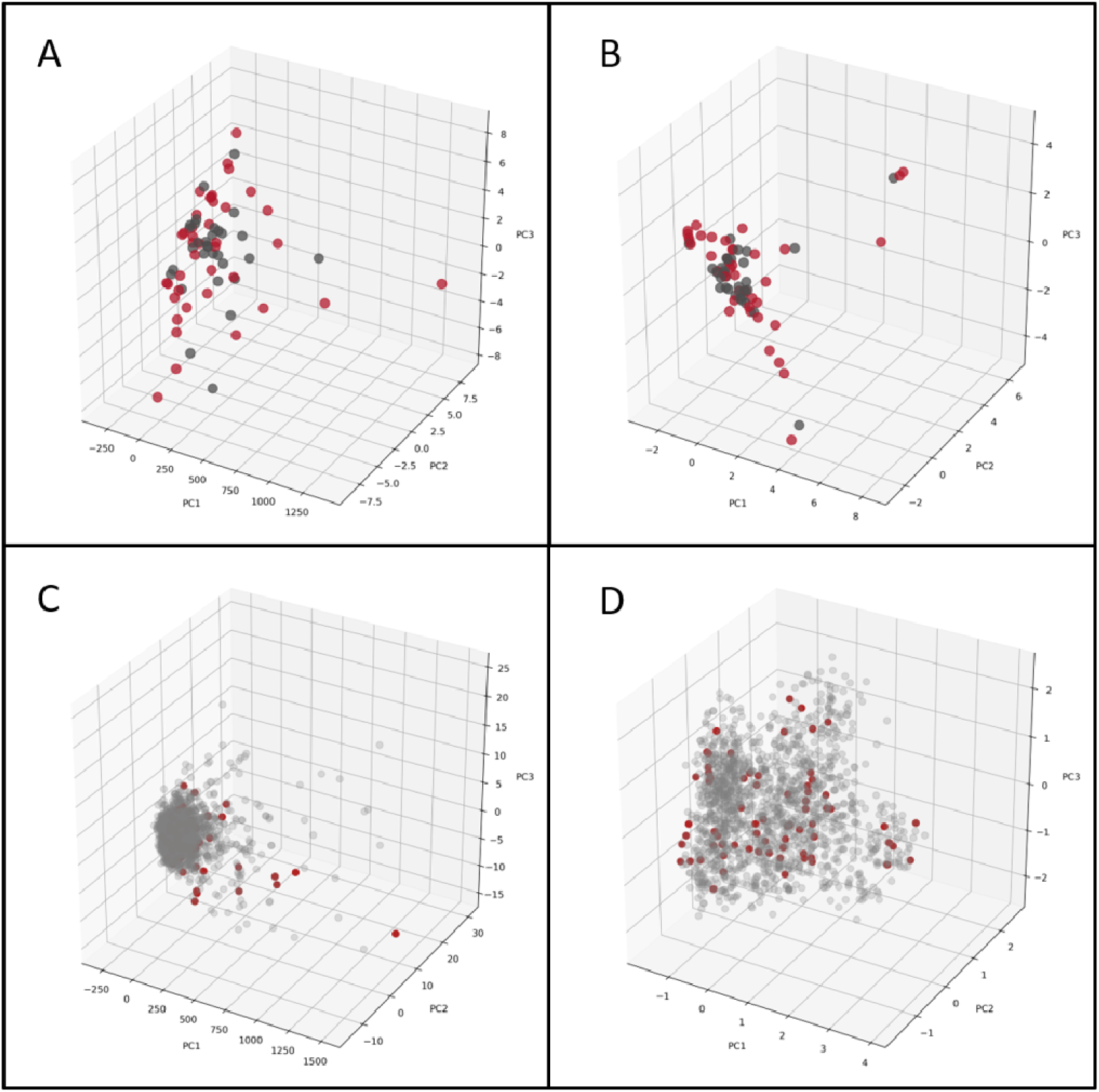
PCA of the SARS-CoV-2 set with Molecular Descriptors (A), and ECFP6 (B). Red Spheres – Active, Grey Spheres – Inactive. PCA of SARS-CoV-2 set and Prestwick Chemical Library (PwCL) with molecular descriptors (C), and ECFP6 (D). Red Spheres – SARS-CoV-2, Grey Spheres – PwCL

**Figure 2.**
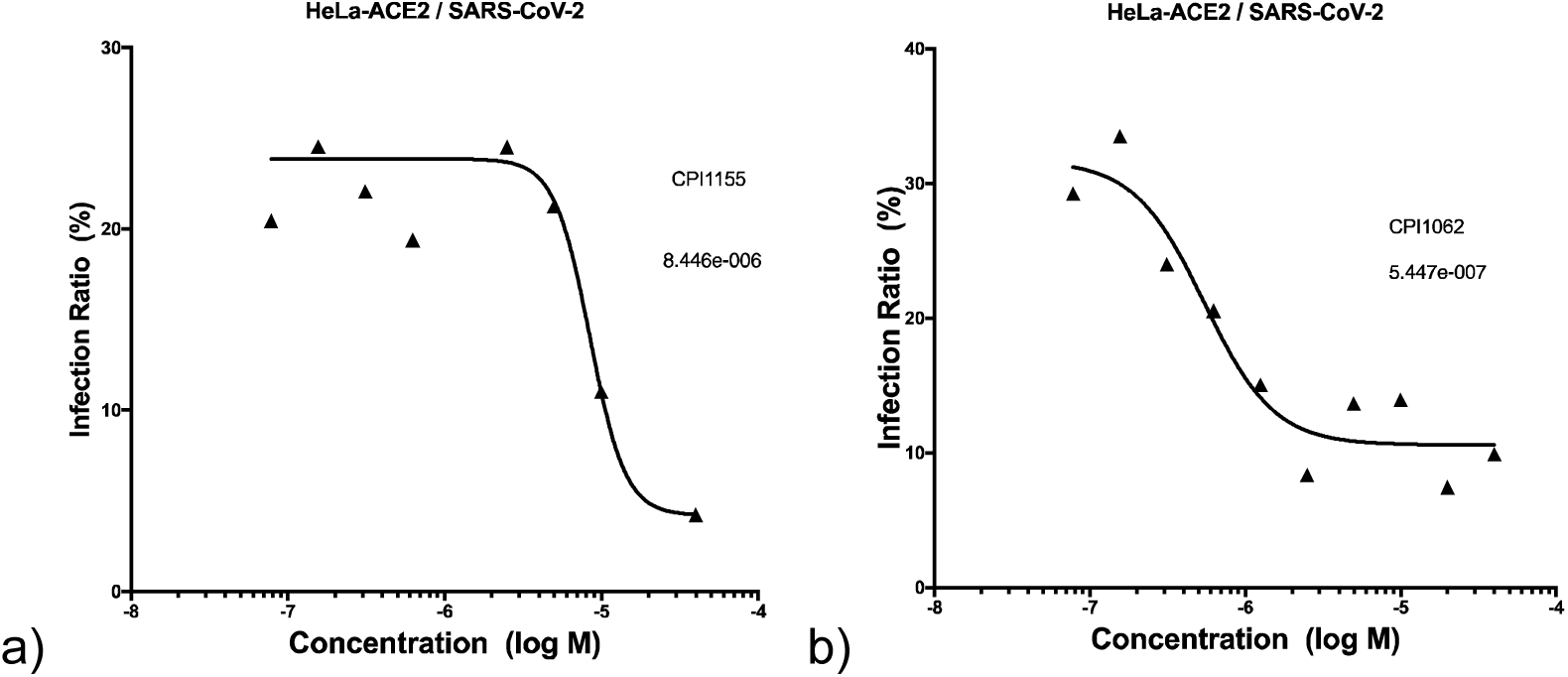
Preliminary dose response curves for a) CPI1155 and b) CPI1062.

The molecules of the dataset do not have a common scaffold, but there are several common structural features that occur in active/inactive molecules that can be highlighted, such as tertiary amines and aliphatic chains in active molecules and phenyl rings and peptide molecule features in inactive molecules (Figure S3). These most common active features appear in chloroquine, tripanarol and tilorone, while the inactive features appear in darunavir, amprenavir and ritonavir (Figure 3). The lack of common scaffolds and features that appears in more than 30% of the active or inactive molecules shows how different and diverse the active molecules are, which turn classification models for these molecules into a relatively difficult task.

**Figure 3.**
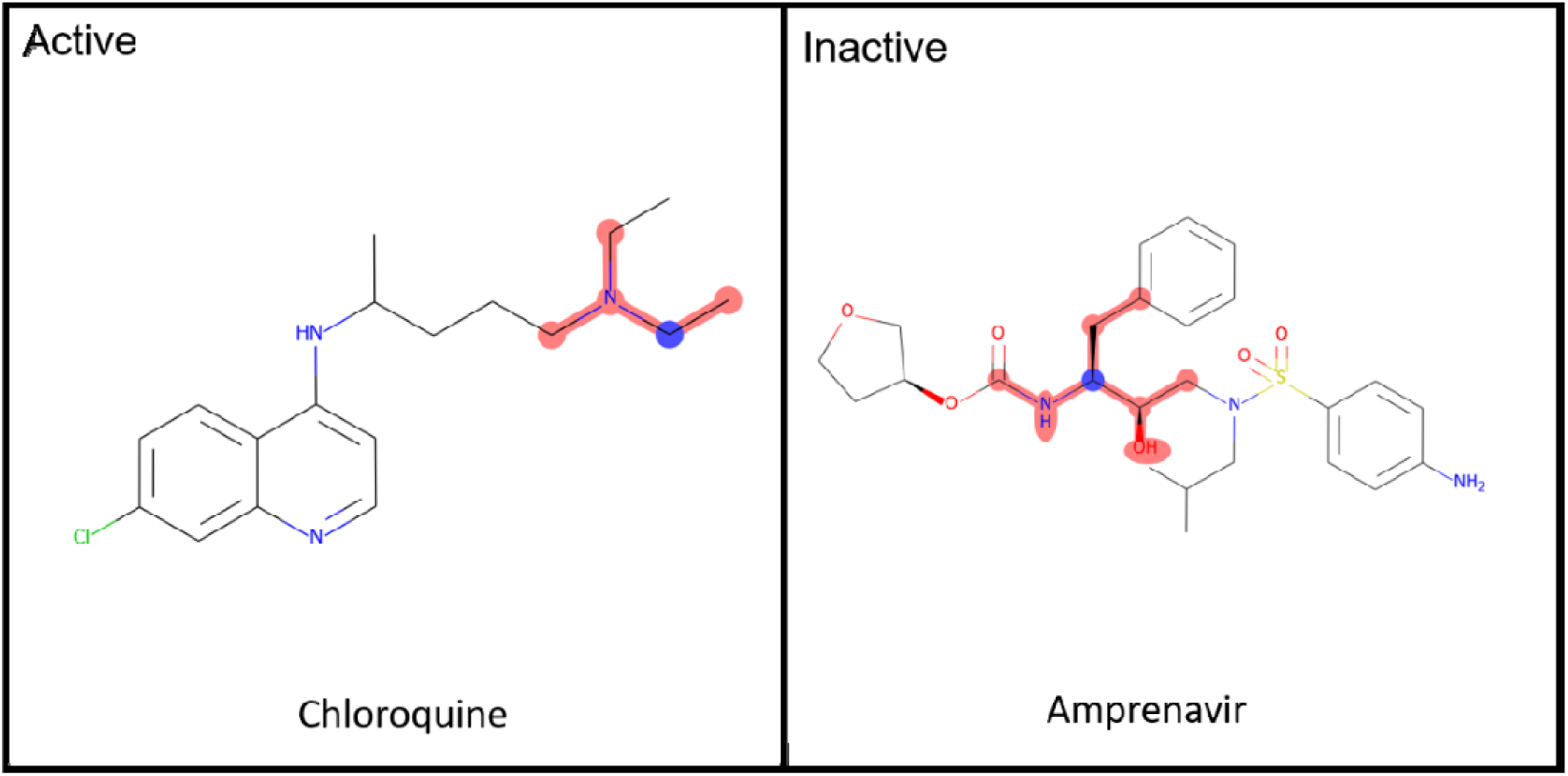
Common Active/Inactive structure features of the SARS-CoV-2 dataset

The performance of a predictive model is highly dependent on the curation and data used. One of the main problems that comes from building models with biological data from different laboratories is data reproducibility and assay standardization^40^. Cell based assays of viral infections have many parameters that can affect the compound potency, e.g., cell lines, MOI, assay readout^41^. From all inhibition assays for SARS-CoV-2 collated to date, most studies use MOI of 0.01-0.05 (73% of data), different Vero cell lines (77% of data) and qRT-PCR (60% of data), however there is no clear definition of compound addition time post infection (Figure S1).

Besides this, even assays with the same or similar conditions have differences in ‘control’ compounds such as chloroquine or remdesivir, showing a lack of data reproducibility between laboratories, which can impact model building. If we keep only studies with the most in common there is not enough data to build a model, while merging all studies will have problems of different assay parameters. It was shown that for Ebola infections in VeroE6 cells the change in the compound potency at different time post infections are lower when using MOI of 0.01-0.1 therefore, merging different assays with the same cell line and low MOI is a good choice to avoid data inconsistency^41^.

It should be noted that most of the *in vitro* data collated to date uses Vero or Vero E6 cells for inhibition assays. Although these cells lines have high ACE2 expression levels, they lack a TMPRSS2 gene. Priming of viral S proteins can occur with the host cell protease TMPRSS2 and Cathepsin L and is essential for SARS-CoV-2 entry^42,43^. Therefore, inhibition assays with cells that do not express TMPRSS2 should be avoided as they might miss compounds that could inhibit the protein and instead find compounds that prevent virus entry by inhibiting only Cathepsin L. In order to avoid these problems with the TMPRSS2 and Cathepsin L gene, cell lines like Calu-3 or modified Vero cell lines should be used instead.^44^

From the 7 compounds prioritized for testing in our laboratory using the machine learning model, CPI1155 and CPI1062 showed antiviral activity against SARS-CoV-2 infections in HeLa-ACE2 cells. Like Vero cells, HeLa does not express TMPRSS2, therefore compounds might need to be be retested in different cell lines to see whether or not the expression of TMPRSS2 affects compound activity.^45^

As new data is continually being published the machine learning models can be updated to increase performance in terms of both training and external test set validation. The very latest model for SARS-CoV-2 is available at www.assaycentral.org. In the meantime, we have shown these models perform well with internal cross validation, external validation as well as prospective prediction, enabling us to find additional active molecules. These models should be used to prioritize compounds which have both a high prediction score and reliability as described herein. This will be expected to return more reliable predictions that together with drug discovery expertise can help prioritize compounds in future for *in vitro* testing.

## Acknowledgements

We would like to kindly acknowledge Dr. Nancy Baker and Ms. Natasha Baker for their help in collating recently SARS-CoV-2 published data. We also thank Biovia for supplying Discovery Studio. Per Subcontract: “This material is based upon work supported by the Defense Advanced Research Projects Agency (DARPA) under Contract No. HR001119C0108.” Per DISTAR Form: “The views, opinions, and/or findings expressed are those of the author(s) and should not be interpreted as representing the official views or policies of the Department of Defense or the U.S. Government.”

## Funding

We kindly acknowledge NIH funding: R44GM122196-02A1 from NIGMS (PI – Sean Ekins) and support from DARPA (HR0011-19-C-0108; PI: P. Madrid) is gratefully acknowledged. Distribution Statement “A” (Approved for Public Release, Distribution Unlimited). The views, opinions, and/or findings expressed are those of the author and should not be interpreted as representing the official views or policies of the Department of Defense or the U.S. Government. FAPESP funding: 2019/25407-2 (PI – Glaucius Oliva).

## Conflicts of interest

SE is CEO and owner of Collaborations Pharmaceuticals, Inc. DHF, KMZ, TRL, AP are employees of Collaborations Pharmaceuticals, Inc.

