## Supplementary material for "Machine Learning Models Identify Inhibitors of SARS-CoV-2": supplmental data

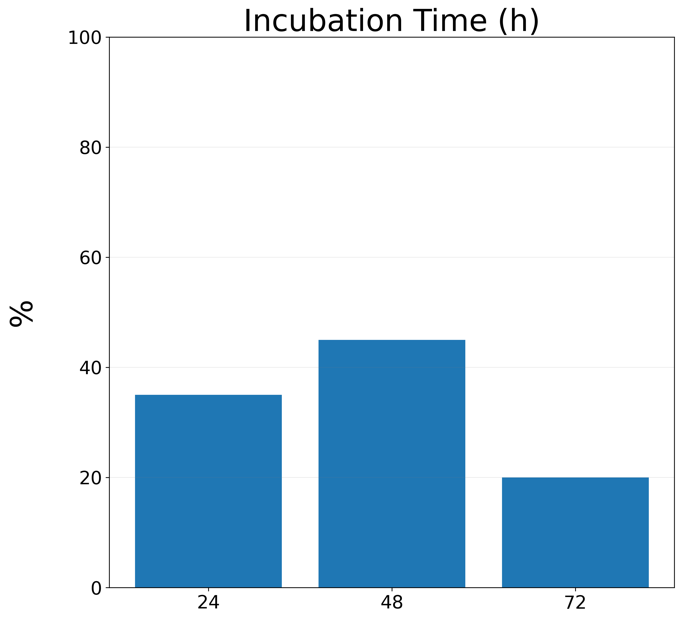

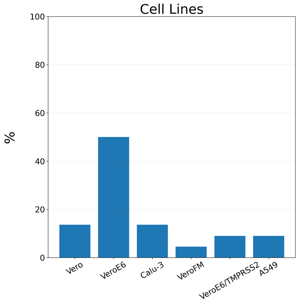

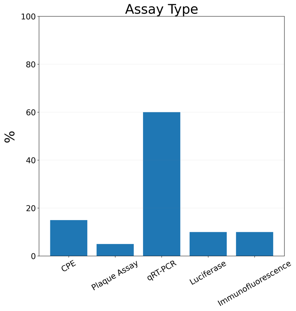

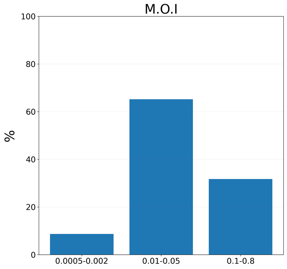

Figure S1 – Percentage of different cell-based assay parameters in the data collated.

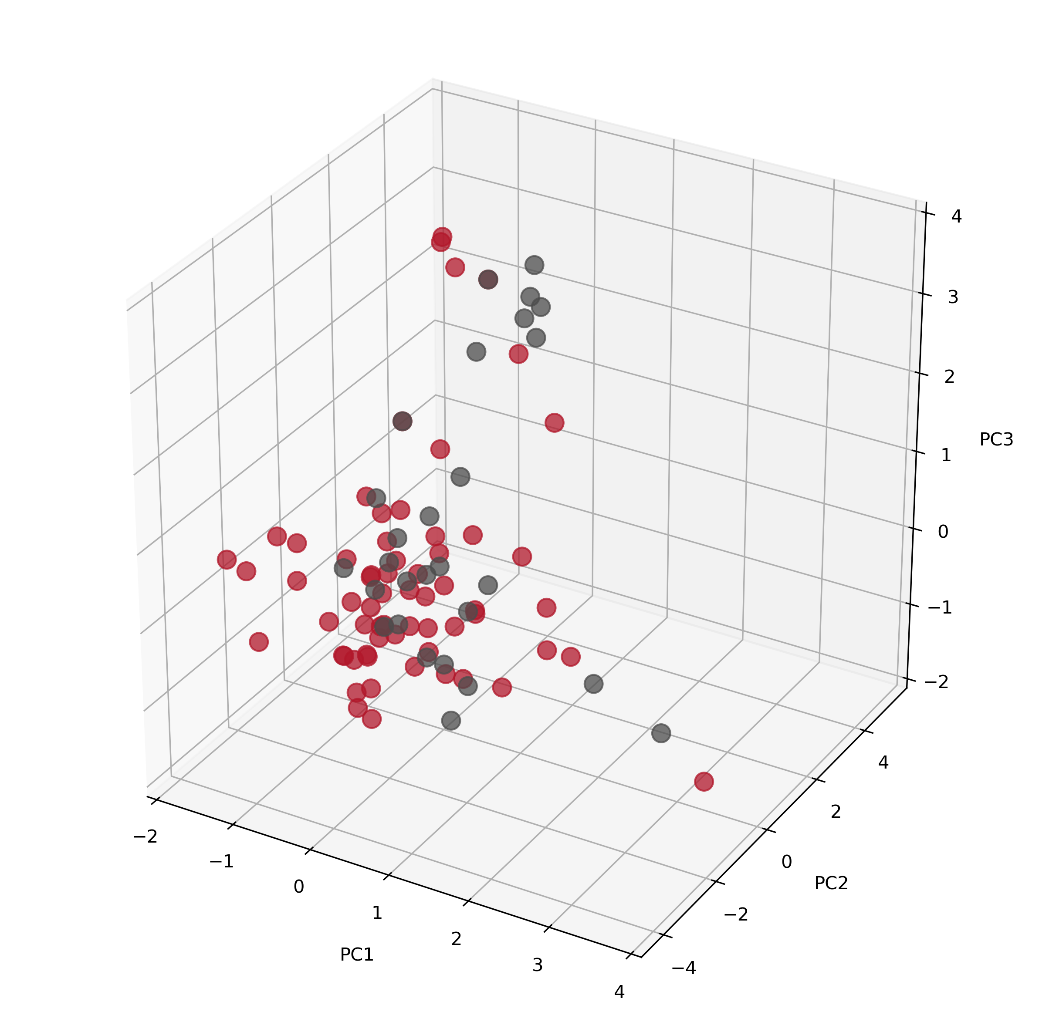

Figure S2 – PCA of test set (gray) and training set (red).

**A**
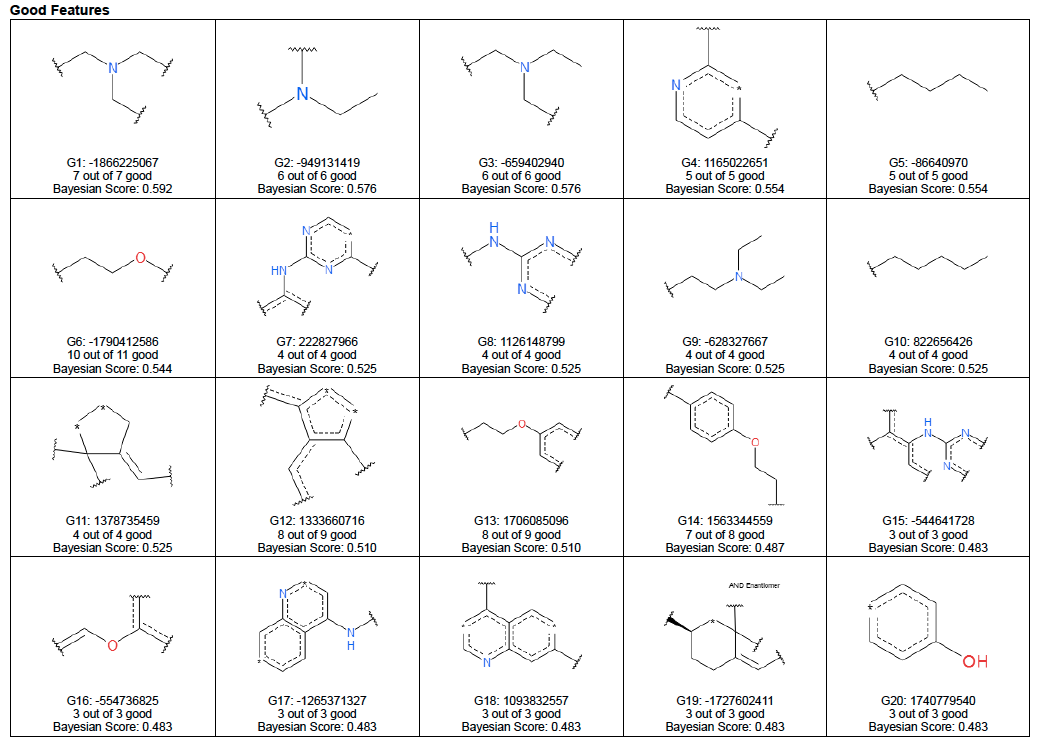

**B**
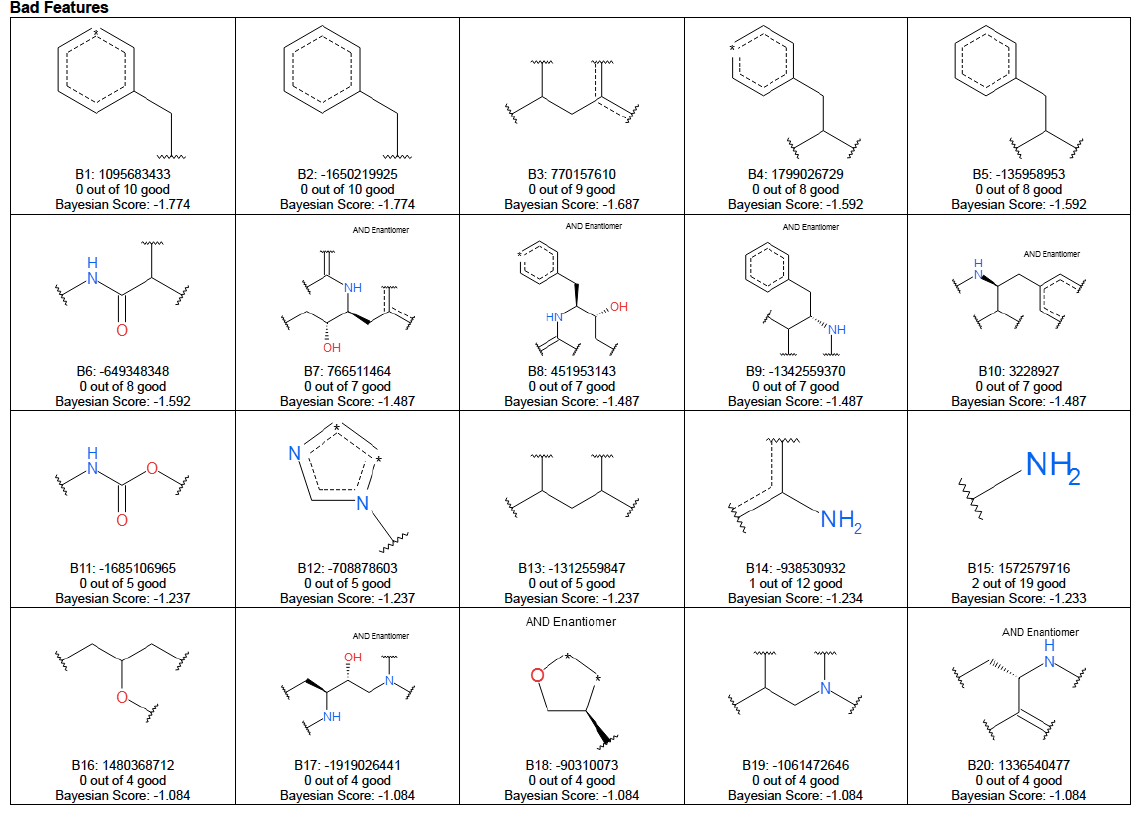

Figure S3. Good and bad molecular features. In order to evaluate the good and bad features of the training data a Bayesian Model with ECFP6 was built with the Discovery Studio 4.0 (BIOVIA) using -fold cross validation and default parameters. Model Statistics: ROC: 0.736, Recall: 0.891, Specificity: 0.894, Accuracy: 0.89, MCC: 0.7849, Precision:0.89, F1:0.89.

**Supplemental Methods**

**Reliability Domain Criteria Definition and Scores**

1) Does the compound fall within the general coverage of the training set?

The distance between test compound and the major central point of the training set are computed based on ECFP6 fingerprints. If the compound is within the maximal radius of the major central space, it falls within the general coverage of the training set and gets a score of 1. If not, it gets a score of 0.

- Yes (1): Within the general coverage of the training set
- No (0): Outside the general coverage of the training set

2) Does it fall within reliable distance from the closet cluster(s)? (higher priority)

The distances between the test compound and all the centroids from the training set clusters are computed. Identify the clusters to which the test compound belongs based on whether it falls within the maximal radii of the individual clusters (note that there can be multiple clusters in which the test compound falls). For the clusters that it belongs to, determine the reliability index of -1, 0 or 1. In case of many clusters identified, choose a single one by computing the reliability indexes for those clusters and selecting the one with the highest reliability index. If there are no clusters, choose the nearest cluster based on the simple distance calculation.

- Low (-1): the compound falls outside the maximal radius of the nearest cluster.
- Moderate (0): Near the far end of the cluster since the compound falls between 75^th^ percentile and the maximal radius of the nearest cluster.
- High (1): Well within the coverage of the cluster since the compound falls within 75^th^ percentile of the nearest cluster.

3) Is the closest cluster dense enough? (higher priority)

It is an indication of the sparsity/density of a cluster. The strength of cluster density for all the internal clusters are represented by values of -1, 0, 1.

- Sparse cluster (-1): It is a sparse cluster because it contains less than 60% of the cluster members that an average cluster should contain.
- Moderately crowded cluster (0): It is a decent cluster because it contains between 30% and 60% of the cluster members that an average cluster should contain.
- High density cluster (1): It is a good cluster because it contains more than 30% of the cluster members that an average cluster should contain.

4) Is the closest cluster(s) well within the chemical space?

It assesses how far the cluster is from the major central point. The strength of whether a cluster belongs in the general chemical coverage of the training are represented by values of -1, 0, 1.

- Far out in the chemical space (-1): This cluster is somewhat outside the chemical coverage of the training set because its centroid distance to the major central point is more than 2 standard deviations among all the distances (absolute zscore >2)
- Moderately within the chemical space (0): This cluster is moderately within the entire chemical coverage of the training set because its centroid distance to the major central point is between one and two standard deviations (1 < absolute zscore < 2)
- Well within the chemical space (1): This cluster is well within the entire chemical coverage of the training set because its centroid distance to the major central point is less than i.e. one standard deviation (absolute zscore < 1)

**Table S1 –** Training Data

| Name | EC50 (µM) | Reference |
| --- | --- | --- |
| Ribavirin | 109.5 | 1 |
| Penciclovir | 95.96 | 1 |
| Favipiravir | 61.88 | 1 |
| Remdesivir | 0.77 | 1 |
| Hexachlorophene | 0.9 | 2 |
| Oxyclozanide | 3.71 | 2 |
| Niclosamide | 0.28 | 2 |
| Pyronaridine tetraphosphate | 31.75 | 2 |
| Berbamine hydrochloride | 7.87 | 2 |
| Penfluridol | 5.01 | 2 |
| Camostat | 50 | 2 |
| Cepharanthine | 4.47 | 2 |
| Tetrandrine | 3 | 2 |
| Mequitazine | 7.28 | 2 |
| Thioridazine hydrochloride | 6.69 | 2 |
| Phenazopyridine | 28 | 2 |
| Loperamide | 9.27 | 2 |
| Perhexiline maleate | 6.38 | 2 |
| Ebastine | 6.92 | 2 |
| Ivacaftor | 6.57 | 2 |
| Atazanavir | 50 | 2 |
| Isopomiferin | 4.51 | 2 |
| Lanatoside C | 50 | 2 |
| Proscillaridin | 2.04 | 2 |
| Osajin | 3.87 | 2 |
| Ciclesonide | 4.33 | 2 |
| Lopinavir | 11.8 | 2 |
| Bazedoxifene | 3.44 | 2 |
| LDK378 | 2.86 | 2 |
| Eltrombopag | 8.27 | 2 |
| Droloxifene | 6.6 | 2 |
| Gilteritinib | 6.76 | 2 |
| Osimertinib mesylate | 3.26 | 2 |
| Pralatrexate | 50 | 2 |
| Cyclosporine | 5.82 | 2 |
| Dronedarone Hcl | 3.92 | 2 |
| Hydroxyprogesterone caproate | 6.3 | 2 |
| Lusutrombopag | 3.78 | 2 |
| Anidulafungin | 4.64 | 2 |
| Clomiphene Citrate | 5.36 | 2 |
| Tilorone | 4.09 | 2 |
| Abemaciclib | 6.62 | 2 |
| Mefloquine Hydrochloride | 5.9 | 2 |
| Toremifene Citrate | 6.36 | 2 |
| Triparanol | 6.72 | 2 |
| Amodiaquine Dihydrochloride | 5.23 | 2 |
| Cinanserin | 20.61 | 3 |
| N3 | 16.77 | 3 |
| Hydroxychloroquine | 4.06 | 4 |
| Chloroquine | 2.07 | 4 |
| Fluphenazine Dihydrochloride | 8.98 | 5 |
| Chlorpromazine Hydrochloride | 4.03 | 5 |
| Clomipramine Hydrochloride | 7.59 | 5 |
| Fluspirilene | 5.32 | 5 |
| Benztropine Mesylate | 17.79 | 5 |
| Gemcitabine Hydrochloride | 50 | 5 |
| Terconazole Vetranal | 16.14 | 5 |
| Imatinib Mesylate | 5.32 | 5 |
| Promethazine Hydrochloride | 10.44 | 5 |
| Anisomycin | 50 | 5 |
| Emetine Dihydrochloride Hydrate | 50 | 5 |
| Tamoxifen Citrate | 8.98 | 5 |
| Thiethylperazine Maleate | 8.02 | 5 |

**Table S2 –** External Test Data

| **Name** | **EC50 (µM)** | **Reference** |
| --- | --- | --- |
| Ivermectin | 2.00 | 6 |
| Galidesivir | 100 | 7 |
| Fludabarine | 100 | 7 |
| Baloxivir | 100 | 7 |
| 4'-Azidocytidine | 100 | 7 |
| Homoharringtonine | 3.12 | 7 |
| Dalbavancin | 100 | 7 |
| Oritavancin | 100 | 7 |
| Tenofovir | 100 | 7 |
| Oseltamivir | 100 | 7 |
| Apilimod | 0.02 | 8 |
| Baicalein | 1.7 | 9 |
| β−D−N4-hydroxycytidine | 0.3 | 10 |
| Sulfadoxine | 35.3 | 11 |
| Clemizole hydrochloride | 24 | 11 |
| Dolutegravir | 22 | 11 |
| Omeprazole | 17 | 11 |
| Azithromycine | 2.12 | 11 |
| Alprostadil | 5.4 | 11 |
| Arbidol | 10.7 | 11 |
| Indomethacin | 1 | 12 |
| Saquinavir | 8.62 | 13 |
| Indinavir | 59.13 | 13 |
| Darunavir | 46.4 | 13 |
| Amprenavir | 31.31 | 13 |
| Ritonavir | 8.62 | 13 |
| Nelfinavir | 1.13 | 13 |
| Tipranavir | 13.34 | 13 |
| Boceprevir | 1.9 | 14 |
| GC376 | 3.37 | 14 |
